# Global heterogeneity of latitudinal patterns in herbivory between native and exotic plants

**DOI:** 10.1101/2024.01.24.576872

**Authors:** Yaolin Guo, Madalin Parepa, Hui Wang, Min Wang, Jihua Wu, Bo Li, Rui-Ting Ju, Oliver Bossdorf

## Abstract

Latitudinal gradient of herbivory that differs between native and exotic plants has been an open issue. It is expected that a latitudinal gradient of herbivory will be evident for native plants; this gradient in exotic plants may lag behind native plants in terms of co-evolution with local abiotic and biotic factors in introduced area. Our study aims to elucidate the difference in latitudinal gradient of herbivory between native and exotic plants globally, while exploring the underlying mechanisms driving the latitudinal gradient of herbivory with biotic and abiotic factors. To achieve this, we analyzed herbivory data from 94 studies and calculated 862 herbivory effect sizes (*z_r_*) to quantitatively characterize the intensity of latitudinal herbivory gradient. For each herbivory data, we matched the corresponding plant identity (native/exotic), herbivore feeding guilds and climate data to reveal the contribution of these factors to *z_r_*. Our findings demonstrate a significant decrease in the latitudinal gradients of with increasing latitude for native plants, a pattern not observed for exotic plants. The heterogeneity in latitudinal gradients of herbivory between native and exotic plants is mediated by herbivore feeding guilds. There is a significant influence of climatic factors on the latitudinal gradient of herbivory for native plants, but not for exotic plants. Overall, our results underscore a general heterogeneity in global macroecological herbivory patterns between native and exotic plants, and highlight the role of biotic and abiotic factors in explaining these global-scale differences.

## Introduction

The latitudinal biotic interaction hypothesis (LBIH) posits that biotic interactions are more intense at lower latitudes compared to higher latitudes (Zvereva & Kozlov, 2021). This hypothesis has been increasingly acknowledged as a crucial explanation for the latitudinal diversity gradient hypothesis (LDGH) (Kinlock et al., 2018). Herbivory, a major component of biotic interactions in terrestrial ecosystems (Schowalter et al., 1986; Schmitz, 2008), has consistently intrigued biogeographers, spurring research into its latitudinal patterns (Anstett et al., 2016). Numerous studies have indicated that herbivory decreases with increasing latitude (e.g., Coley & Aide 1991; Salgado & Pennings 2005; Więski & Pennings 2014). However, Moles et al. (2011) noted in their meta-analysis that merely 37% of studies corroborated the LBIH, evidencing increased herbivory rates at lower latitudes. The underlying causes of these heterogeneous findings are multifaceted, including restricted latitudinal ranges examined and comparisons among different species across latitudes (Schemske et al., 2019; Moles et al., 2011). Particularly, the scarcity of comparative studies on plant identities (native versus exotic plants) within existing research on latitudinal herbivory gradients may also contribute to these inconsistencies.

Large-scale spatial heterogeneity in latitudinal herbivory gradients has, contrasting between native and exotic plants, is observable (Bezemer et al., 2014). A previous field study revealed non-parallel latitudinal gradients in herbivory and related traits between native and exotic lineages of *Phragmites australis* (Cronin et al., 2015). To extend these findings, Bhattarai et al. (2017) performed a common garden experiment within the same research framework to demonstrate the heritability of the heterogeneity in latitudinal herbivory gradients between native and exotic plants. Although these findings are derived from a single plant species at the subspecific level (i.e., two lineages of *Phragmites australis*), the authors of this series of papers believed that the generalizability of non-parallel latitudinal gradients between native and exotic plants across various ecosystems, transcending distinctions in plant taxa at the species, genus, or higher taxonomic levels. More recently, several empirical studies have highlighted the relevance and implications of this phenomenon on plant invasions (Zhang et al., 2021; Guo et al., 2023); however, a comprehensive systematic review explicitly addressing the non-parallel latitudinal gradient hypothesis (NLGH) is still markedly lacking.

Exotic plants, having escaped from specialized natural enemies in their introduced ranges compared to native plants—a phenomenon known as ‘enemy release’— consequently experience differing herbivory pressures (Liu & Stiling, 2006; Chun et al., 2010; Xu et al., 2021). A systematic meta-analysis indicates that herbivory on exotic plants is generally less intense than on native plants, and interestingly, the intensity of natural enemy release did not show significant differences across latitude (Xu et al., 2021). However, in their introduced range, where exotic plants are exposed to attack by a broad spectrum of new generalist herbivores, it may respond specifically based on different herbivore feeding guild, although the formation of such complex mechanisms is a long-term co-evolutionary process (Joshi & Vrieling, 2005; Parker et al., 2006). Therefore, the variance in plant response to herbivores might explain be a key factor explaining the different latitudinal gradients in herbivory observed between native and exotic plant.

Climatic factors play a key role in influencing various plant palatability traits and mediating pressure from natural enemies, thereby shaping the latitudinal gradient of herbivory in native plants (Abdala-Roberts et al., 2016). Conversely, exotic plants, which often lacking pre-adaptation to novel environments they encountered across extensive latitudinal ranges, may not initially exhibit similar latitudinal gradients in herbivory and palatability as compared to native plants (Bezemer et al., 2014). Supporting this, numerous studies, such as Liu et al. (2021) have demonstrated that herbivory-related defense syndromes in the widely invasive species *Alternanthera philoxeroides* are significantly responsive to climatic variation in its native range, but not in its introduced range. Similarly, Xiao et al. (2020) found that in the native range of the tallow tree (*Triadica sebifera*), there is a decrease in allocation to palatability traits such as tannins and flavonoids with increasing latitude – a pattern that is absent in its invasive range.

Here, we performed a comprehensive meta-analysis to synthesize empirical research on the different latitudinal gradients in herbivory between native and exotic plants. Our primary goal was to quantify and compare general latitudinal herbivory gradients in native versus exotic plants. Secondly, we sought to explore the underlying complex influence of biotic and abiotic factors on differences in latitudinal herbivory gradient between native and exotic plants. Third, we aimed to investigate the potential for bias that defined as a systematic error in results that favor one outcome over others. Based on previous studies, we hypothesized that: (1) the latitudinal herbivory pattern in native plants decreases with increasing latitude, whereas exotic plants exhibit no significant latitudinal pattern; and (2) this spatial heterogeneity of herbivory arises from ‘response differences’ of native and exotic plants to abiotic and biotic factors across wide ranges. We tested these hypotheses by conducting a meta-analysis of the findings from 96 publications that report on latitudinal variations in herbivory.

## Methods

### Literature search

We searched Web of Science for studies on latitudinal variation in herbivory that were published before 2023. We used the search string ‘invasi* OR exotic OR alien OR introd* OR nativ* OR naturalized plant AND latit* OR geograph* OR biogeograph* AND herbiv* OR leaf area damage OR leaf damage OR herbivory ratio’, which resulted in a total of 3285 papers. Subsequently, we meticulously screened the titles, abstracts and main texts of these papers. Studies were included if they satisfy the following criteria: (1) the study must report herbivory for all individuals or sites included in the study; (2) the study must report the plant species, and we rejected studies that compared the herbivory of different plant species; (3) the data had to be derived from at least two different latitudes; (4) for studies with only two latitudes, it is necessary to report the mean and variance values; (5) the study must report precise latitude and longitude details for all observation sites; (6) the major effect must be latitudinal variation, and we have excluded data resulting from multiple treatment interactions, such as latitude vs. elevation, latitude vs. predator exclusion, and latitude vs. other treatments; additionally (7) we also have excluded several studies that collected plants and conducted experiments at different transcontinental locations.

### Data extraction

A total of 96 published papers fulfilled the inclusion criteria specified above. We used PlotDigtizer (version 3.2.1) to extract data on latitudinal herbivory from the figures in these papers. We identified three primary indicators of herbivory for our analysis: leaf damage area (m^2^), leaf damage rate (%) and herbivore abundance. These parameters are widely recognized as key indicators for assessing the intensity of herbivory (Abdala-Roberts et al., 2016; Moreira et al., 2018). Subsequently, we recorded the latitude and longitude corresponding to each herbivory data. In cases where studies reported only two latitudinal points, we extracted the standard errors (SE) directly or calculated them using the provided standard deviations (SD). To avoid potential bias arising from inter-annual variation in studies spanning multiple years, we extracted data for each year separately. Overall, our analysis included 862 cases.

For each herbivory data, we recorded the corresponding plant species name and its identity as either native or exotic plant. In instances where the plant identity was not explicitly detailed in the study, we consulted the Plants of the World Online database (https://powo.science.kew.org/) for accurate classification and clarification.

For each herbivory data, we matched climatic factors for latitudinal locations with corresponding climatic factors, using data from Worldclim (https://www.worldclim.org). Our database incorporated various climatic parameters, including BIO1 (annual mean temperature), BIO4 (temperature seasonality, standard deviation *100), BI05 (max temperature of warmest month), BIO6 (min temperature of coldest month), BIO12 (annual precipitation), BI013 (precipitation of wettest month), BIO14 (precipitation of driest month), BIO15 (precipitation seasonality, coefficient of variation). We included these specific climatic factors in our database as they are widely recognized for their influence on plant traits and growth, and have been extensively utilized in prior research to explore their relationship with herbivory (Abdala-Roberts et al., 2016; Moreira et al., 2018). Additionally, we recorded the observation types (field survey or common garden) for each herbivory data to verify if plants of both identities exhibited potential latitudinal herbivory gradients on different environmental gradients. This because the distinction was crucial as common garden experiments offer a controlled environment gradient compare to field surveys.

We also recorded herbivore feeding guilds for each herbivory data to assess the effects of different herbivore guilds on latitudinal herbivory gradient. Regarding the herbivore feeding guild in each case, we classified it into one of eight categories (such as folivores, defoliators, gallers, grazers, miners, sap-feeders, seed-feeders, stem-feeders) following the classification in Zvereva & Kozlov (2021).

Finally, we recorded the study year, the publishing journal’s name and its impact factor to evaluate the bias. We quantified the visibility of the journals where these studies were published using their five-year impact factors (IF), as obtained from the ISI Web of Sciences. For those publications lacking an IF, we assigned a default value of IF = 0.

### Effect sizes calculation

We used effect sizes (*z_r_*) to indicate the intensity of the latitudinal gradient of herbivory. We calculated Pearson’s coefficient of correlation (*r*) between herbivory and latitudes, and transformed to a normalized effect sizes using Fisher’s *z* transformation (*z_r_*) (Hillebrand, 2004; Kinlock et al., 2018; Zvereva & Kozlov, 2021):

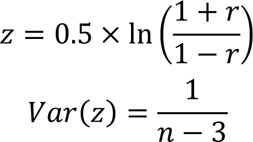

When a study compared herbivory at only two latitudinal sites, we calculated Hedge’s d based on the difference in herbivory between two sites with extreme latitudes, and then converted d to *r* using a web-calculator (www.psychometrica.de/effect_size.html). Hedges ‘d was calculated as:

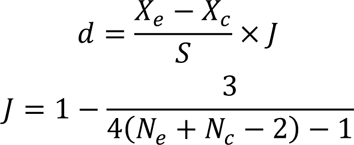

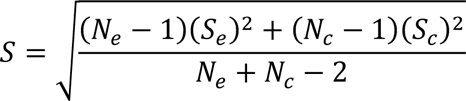

### Meta-analysis

We determined the weighted mean of effect sizes for the entire dataset using the restricted maximum likelihood (REML) method (Cally et al., 2019). Weighted means were calculated by fitting a model without moderating variables (i.e., fixed effects), while incorporating plant species ID as a random effect. This involved separately calculating correlations for the same plant species and taking these interdependencies into account when estimating the overall effect sizes. Subsequently, we designated plant identity as a key moderating variable in our REML models, maintaining plant species ID as a random/group-level effect, to obtain the predictions of effect sizes specific to each plant identity.

To assess the response of plant herbivory to climate, we first calculated the effect sizes (*z_r_*) of climatic factors to represent the intensity of latitudinal gradient in these factors for all studies, using the same method with calculating herbivory effect sizes. Subsequently, we separately fitted climatic effect sizes with herbivory effect sizes of native and exotic plants to test whether greater climatic variability along latitude might lead to more pronounced latitudinal patterns of herbivory. Additionally, we calculated the latitude range (latitude _max_-latitude _min_) and longitude range (longitude _max_ - longitude _min_) for all studies to fit the relationship between both ranges and herbivory effect sizes (Figure S2). This approach is based on the premise that larger latitudinal ranges are indicative of greater climate variability. Furthermore, to reflect the response level of both plant identities to climate gradients, we ran a REML model to analyze the effect of different observation types (field survey or common garden) on latitudinal herbivory gradients. We first ran a full model with fixed observation types as the main moderator for the full dataset, while keeping the plant species ID as a random/group-level effect to obtain the prediction of effect sizes for each plant identity. We then added the plant identity to the model, fixed plant identity, observation types, and their interaction as a moderate variable in the REML models, while keeping the plant species ID as a random effect to derive the prediction of effect sizes for each class of observation types.

To assess the contribution of herbivore feeding guilds to the heterogeneity of latitudinal herbivory gradient between native and exotic plants, we first ran a full model with fixed herbivore feeding guilds as the main moderator for the full dataset, while keeping the plant species ID as a random/group-level effect to obtain the prediction of effect sizes for each plant identity. We then added the plant identity to the model, fixed plant identity, herbivore feeding guilds, and their interaction as a moderate variable in the REML models, while keeping the plant species ID as a random effect to derive the prediction of effect sizes for each class of herbivore feeding guilds. We also fitted a relationship between introduction year and herbivory effect sizes of exotic plants to indicate whether longer introduction time might enhance the strength of latitudinal gradient in herbivory (Figure S4).

Finally, we provided estimates of the heterogeneity present in the dataset. We used the statistic *I*^2^ as an estimate of the proportion of variance in effect sizes due to differences between random effect levels (Viechtbauer, 2010). *I*^2^ is superior to other statistics because it is independent of sample size, easy to interpret, and can be allocated between random effects (Nakagawa et al., 2017).

Our meta-analysis of all REML model fits was implemented using the *metafor* package in R (version 4.2.3) (Egger et al., 1997).

### Assessment of publication bias

Analyzing and accounting for publication and confirmation biases, which are widespread in ecological research (Holman et al., 2015; Jennions et al., 2012; Zvereva & Kozlov, 2019), would help to reach unbiased conclusions despite the fact that some of the published data may be biased. We checked the publication bias using Egger’s test to quantify mapping asymmetry, and checked it by funnel plots (Egger et al., 1997). We also tested the time-lagged bias (Murtaugh, 2002), where the magnitude of the effect sizes decreases over time as more data are collected. Furthermore, we assessed potential sources of publication bias through the correlation between effect sizes and journal impact factor (Jennions & Møller, 2002), which occurs if null or opposite results are more difficult to publish (impact factor from InCites Journal Citation Reports).

## Results

### The entire dataset

We extracted 13869 observed values of herbivory from 94 studies worldwide (Figure 1a; Table S1). Based on these data, we calculated 862 effect sizes of the intensity of latitudinal gradient in herbivory. Among them, 91 cases focused on exotic plants, while 771 cases were measured on native plants. Most cases in our dataset came from studies that were observed in the field (n = 809); 53 cases were derived from common garden experiments (i.e., all raw data observed under the same environment factors). We also obtained effect sizes for 19 climatic factors that matched the above herbivory effect sizes (n = 862).

**Figure 1.**
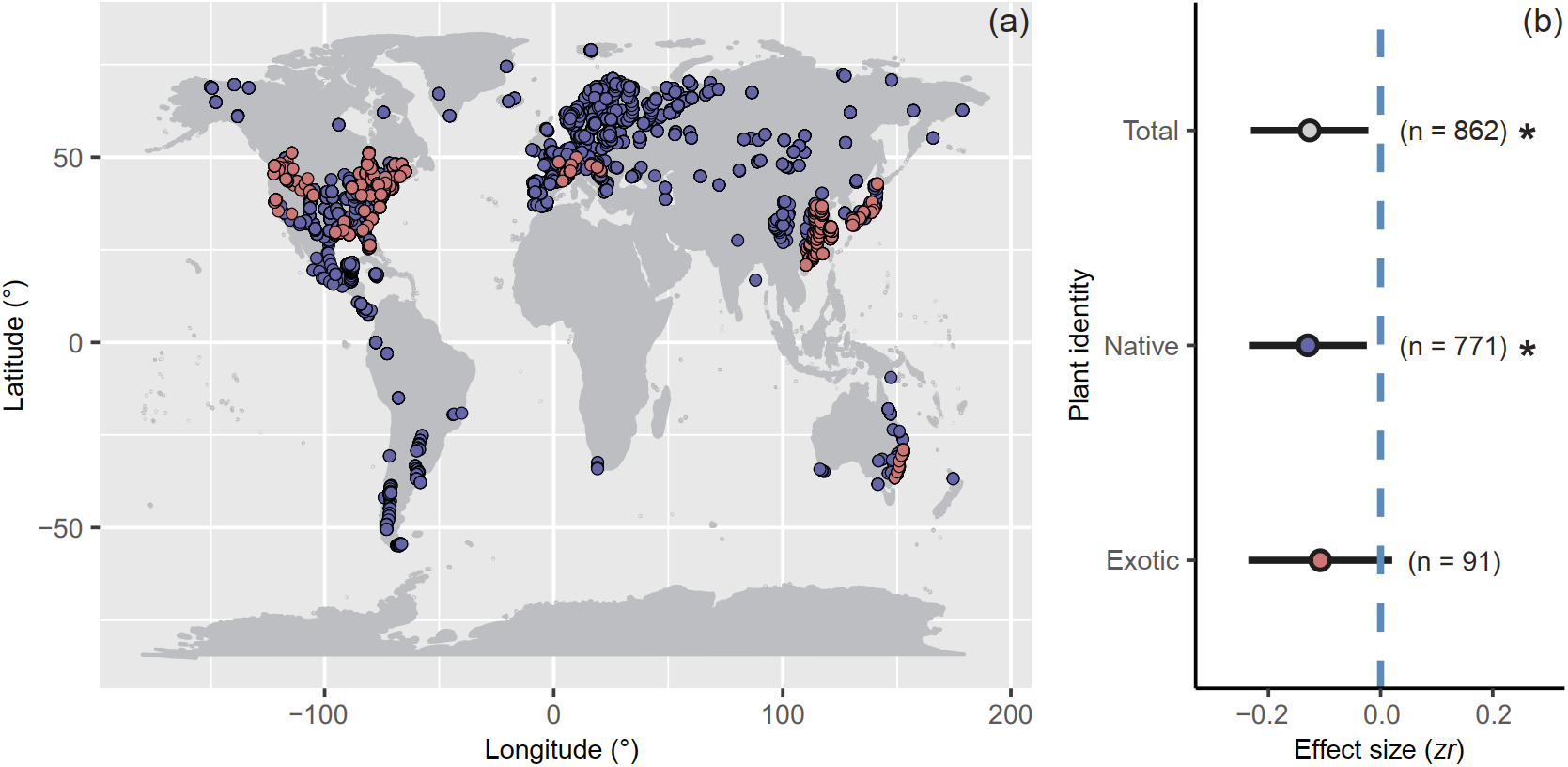
All original observation site for herbivory of exotic and invasive plants (a), the role of plant identities on effect sizes (b). On the panel (a), several studies that only provided latitude without longitude were deleted from the map, but included in the following analysis on effect sizes. On the panel (b), the negative effect sizes indicate a decreased interaction intensity with increased latitude. The estimates and 95% confidence intervals (i.e., horizontal lines) presented here are from restricted maximum likelihood (REML) models. Numbers of effect sizes are shown in parentheses. 95% CI of the linear regression. Statistical significance is indicated as: *, *p* < 0.05; **, *p* < 0.01; ***, *p* < 0.001.

### Effects of the plant identity on latitudinal gradient in herbivory

The overall strength of the latitudinal gradient in herbivory was significantly negative (REML β = −0.127, 95% CIs: −0.231 − −0.022; Figure 1b; Table S1), indicating that the intensity of herbivory decreases with increasing latitude. Plant identity (i.e., native and exotic) significantly influenced the effect sizes (i.e., the intensity of latitudinal gradient in herbivory). The effect sizes of native plants were significantly negative, and similar to the overall effect sizes (REML β = −0.130, 95% CIs: −0.235 − −0.025; Figure 1b; Table S1). In contrast, the effect sizes of exotic plants crossed zero point and were therefore not significant (REML β = −0.108, 95% CIs: −0.236 − 0.021; Figure 1b; Table S1).

### Effects of the herbivore feeding guilds on latitudinal gradient in herbivory

For all dataset, only the effect sizes of all folivores (REML β = −0.127, 95% CIs: −0.238 −-0.016; Figure 2a; Table S1), defoliators (REML β = −0.130, 95% CIs: −0.241 − −0.019; Figure 2a; Table S1), sap-feeders (REML β = −0.171, 95% CIs: −0.301 − −0.040; Figure 2a; Table S1), seed-feeders (REML β = −0.215, 95% CIs: −0.396 − −0.034; Figure 2a; Table S1), and stem-feeders (REML β = −0.349, 95% CIs: −0.515 − −0.182; Figure 2a; Table S1) were significantly negative value. We then distinguished the plant identity to investigate the effect of the herbivore feeding guild on latitudinal gradient in herbivory. For native plants, only the effect sizes of all folivores (REML β = −0.138, 95% CIs: −0.250 − −0.025; Figure 2b; Table S4), defoliators (REML β = −0.148, 95% CIs: −0.261 − −0.035; Figure 2b; Table S4), and stem-feeders (REML β = −0.427, 95% CIs: −0.602 − −0.253; Figure 2b; Table S4) were significantly negative. For exotic plants, only the effect sizes of sap-feeders (REML β = −0.279, 95% CIs: −0.454 − −0.105; Figure 2b; Table S4) and seed-feeders (REML β = −0.512, 95% CIs: −0.857 − −0.168; Figure 2b; Table S4) were significantly negative. There is no evidence to show that the other feeding guilds can influence the latitudinal gradient of herbivory, as the zero point of non-significance was crossed.

**Figure 2.**
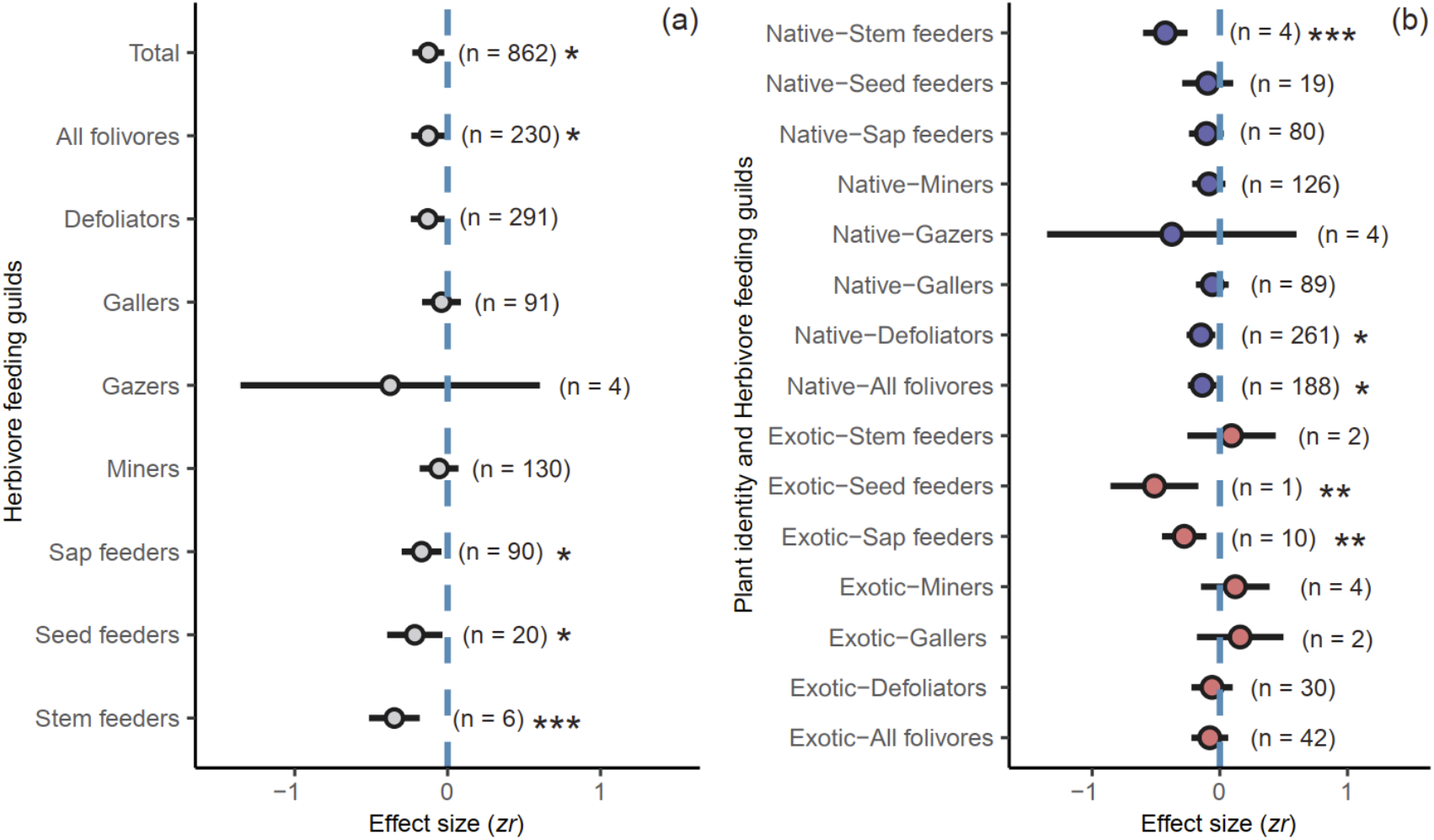
The role of plant identities and herbivore feeding guilds on effect sizes. On panels graph, the negative effect sizes indicate a decreased interaction intensity with increased latitude. The estimates and 95% confidence intervals (i.e., horizontal lines) presented here are from restricted maximum likelihood (REML) models. Numbers of effect sizes are shown in parentheses. Statistical significance is indicated as: *, *p* < 0.05; **, *p* < 0.01; ***, *p* < 0.001.

### Effects of the climatic factors on latitudinal gradient in herbivory

For native plants, the effect sizes for BIO1 (annual mean temperature), BIO12 (annual precipitation), BIO13 (precipitation of wettest month), BIO14 (precipitation of driest month) showed a significant positive correlation with the herbivory effect sizes. Conversely, the effect sizes for BIO4 (temperature seasonality, standard deviation *100) exhibited a significant negative correlation with herbivory effect sizes in native plants (Figure 3). In the case of exotic plants, no significant relationship was found between climate effect sizes and herbivory effect sizes (Figure 3).

**Figure 3.**
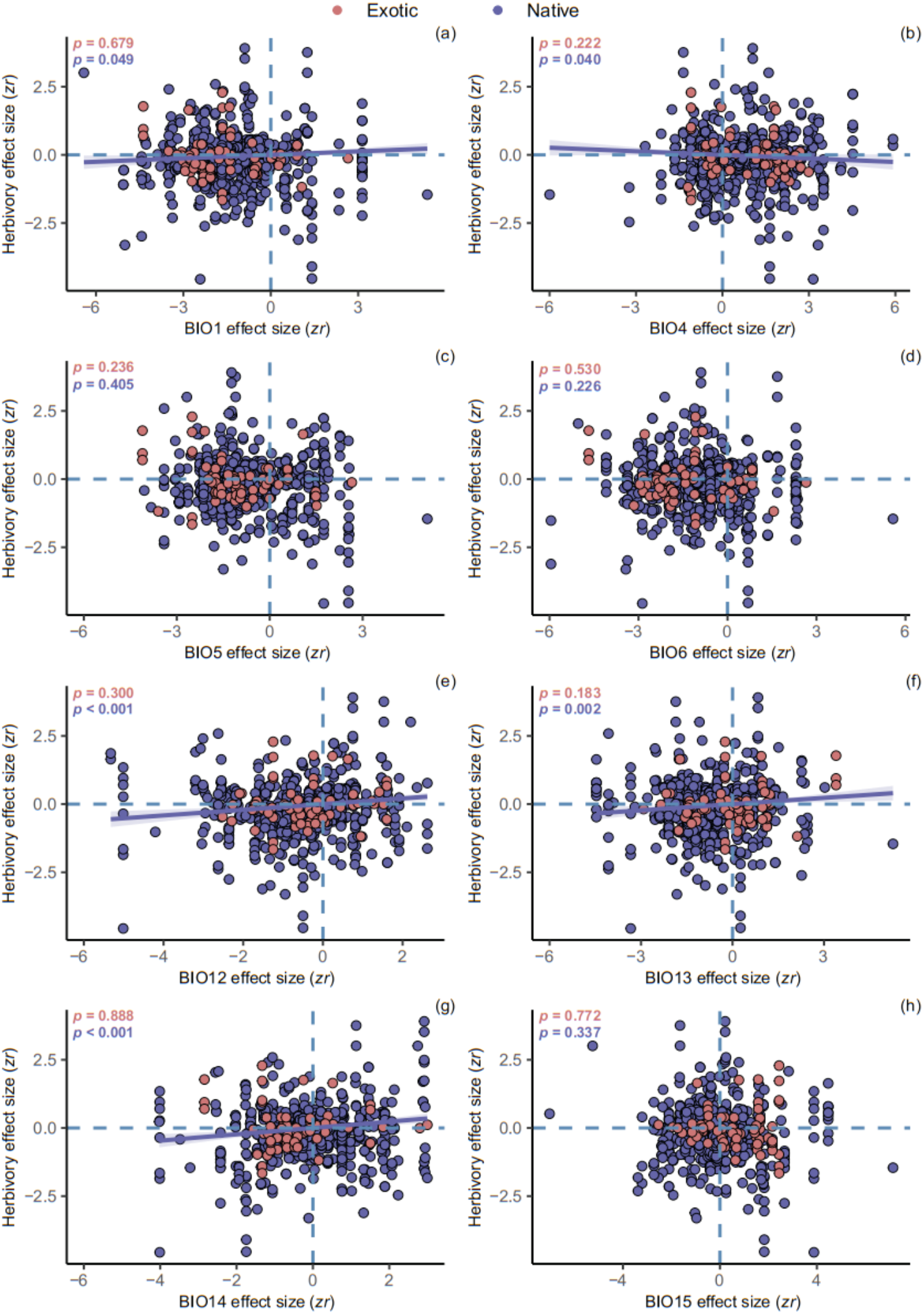
The relationships between effect sizes and main climatic factors. On the graph, BIO1 = Annual Mean Temperature, BIO4 = Temperature Seasonality (i.e., standard deviation *100), BIO5 = Max Temperature of Warmest Month, BIO6 = Min Temperature of Coldest Month, BIO12 = Annual Precipitation, BIO13 = Precipitation of Wettest Month, BIO14 = Precipitation of Driest Month, BIO15 = Precipitation Seasonality (i.e., coefficient of variation). On each panel, *p* < 0.05 indicates a significant relationship. The envelopes represent the 95% CI of the linear regression.

For native plants, the greater the range of study cases (including latitude and longitude), the more significantly the herbivory effect sizes deviate from zero. For exotic plants, a similar relationship did not exist (Figure S1).

The effect sizes observed from the field survey were significantly negative similar to the overall effect sizes (REML β = −0.130, 95% CIs: −0.235 − −0.025; Figure S2a). In contrast, the effect sizes from the common garden crossed the zero point and were, therefore, not significant (REML β = −0.108, 95% CIs: −0.236 − 0.021; Figure S2a). Furthermore, we found that the plant identity (i.e., exotic or native) and the observation types (i.e., field survey or common garden) interacted to influence the intensity of latitudinal gradient in herbivory. The difference in observation types significantly altered the effect sizes for native plants, with significant negative effect sizes observed in the field (REML β = −0.142, 95% CIs: −0.246 − 0.037; Figure S2b) and non-significant effect sizes for native plants observed in the common garden (REML β =-0.033, 95% CIs: 0.160 − 0.095; Figure S2b). Moreover, there is no evidence to show that the observation types can affect exotic plants, as the zero point of non-significance was crossed in the field surveys (REML β = −0.072, 95% CIs: −0.205 − 0.062; Figure S2b) and the common garden (REML β = −0.119, 95% CIs: −0.278 − 0.040; Figure S2b).

### The relationship between introduction year and latitudinal gradient in herbivory

The herbivory effect sizes in exotic plants demonstrated a significant positive trend with increasing duration of introduction (Figure S3). This implies that a longer period of introduction may contribute to the development of a latitudinal pattern of herbivory in exotic plants, similar to the pattern observed in native species.

### Publication bias

The funnel plot of effect sizes was symmetrical, suggesting that some publication bias might be non-existence (Figure 4a; Egger’s test: z = −0.861, *p* = 0.390). Linear regressions show the non-significant relationship between effect sizes and journal impact factor (Figure 4b; t = −0.742, *p* = 0.458) or year of publication (Figure 4c; t = 0.127, *p* = 0.899); thus, we found no evidence that effect sizes dictate the likelihood of publication in high-profile journals, or that effect sizes have diminished as the field has matured.

**Figure 4.**
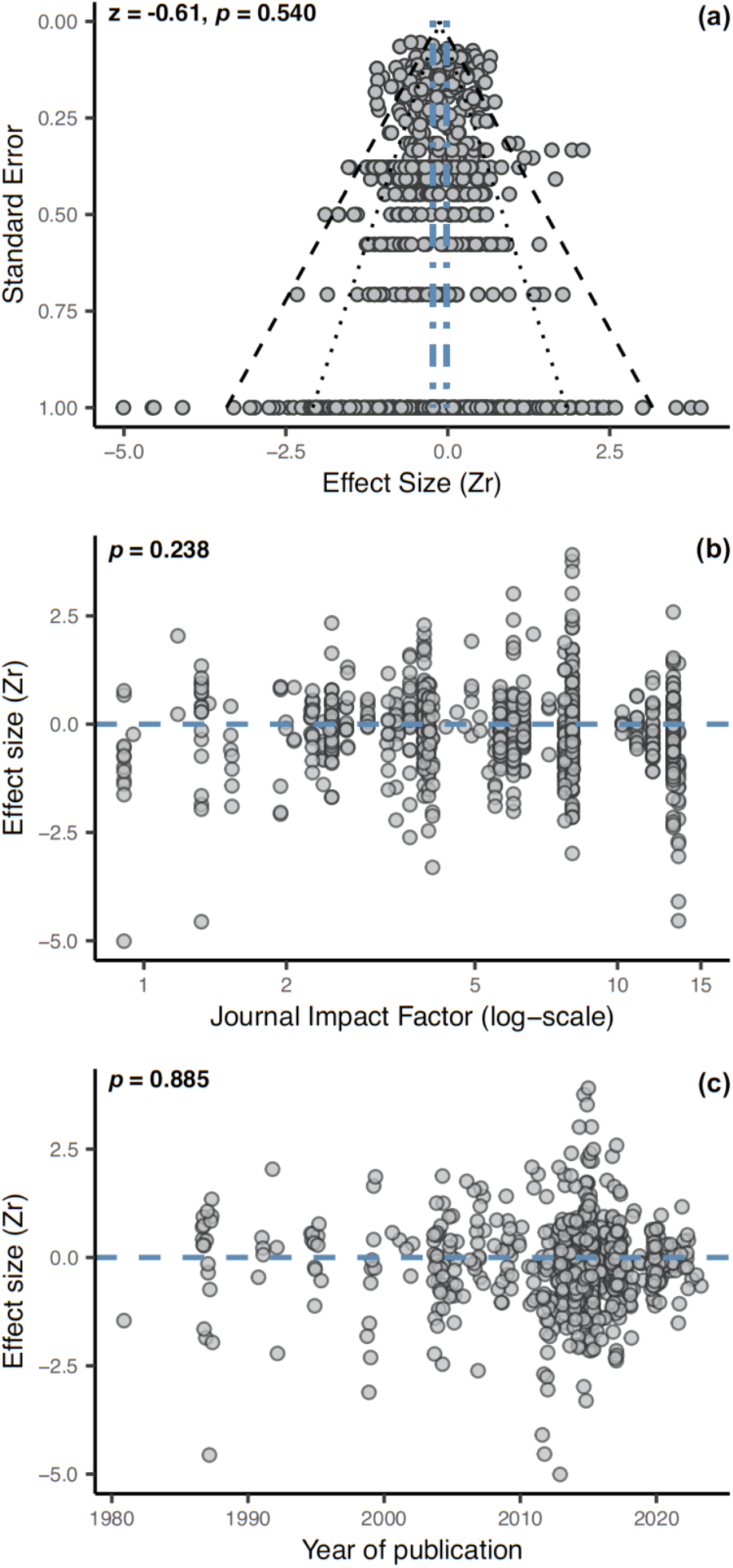
Tests for publication bias in the dataset. On the panel (a), *p* > 0.05 indicates the symmetry of the funnel plot and the inexistence of a publication of bias. All the dashed line was added as visual reference for symmetry. The dark dashed blue lines represent the 95% confidence intervals of the grand mean estimate for all effect sizes, and the black dotted and dashed lines depict the 95% and 99.8% CIs for the dataset. The panel (b) and (c) show the relationship between journal impact factors and effect sizes, and between the year of publication and effect sizes. *p* > 0.05 indicates a non-significant relationship.

## Discussion

In our study, we established an extensive dataset comprising nearly 14,000 observations derived from 861 cases, encompassing over 230 plant species worldwide. Based on this dataset, we reported a robust result of latitudinal patterns in herbivory (Figure 1a). Specifically, we found that a significant difference between native and exotic plants, with native plants exhibiting significant negative latitudinal gradients in herbivory, whereas such patterns were absent in exotic plants (Figure 1b). We then elucidated the potential factors contributing to this difference. Our findings showed that the differential responses to climatic factors and herbivore feeding guilds emerged as essential determinants in shaping the contrasting latitudinal gradients in herbivory of native versus exotic plants (Figure 2; Figure 3). Overall, the significance of our study was revealed a substantial difference in the latitudinal gradient of herbivory between native and exotic plants, thereby providing robust support for the non-parallel latitudinal gradient hypothesis (NLGH).

Our study, based on independent plant species, supports previous study that the intensity of herbivory decreases significantly with increasing latitude (Figure 1b; Zvereva & Kozlov, 2021). Such a result is not similar between native and exotic plants. Specifically, we found a significant negative latitudinal gradient in herbivory of native plants, but not of exotic plants (Figure 1b). Our results suggest that the latitudinal gradient of herbivory is not consistent between native and exotic plants. Thus, we have provided positive evidence in support of the non-parallel latitudinal gradient hypothesis (NLGH) initially proposed by Cronin et al. (2015) and Bhattarai et al. (2017). In addition, we also extended the NLGH that obtained from the study of subspecific taxa (i.e., co-occurring native and invasive lineages of *P. australis*), and increasing its applicability to the biogeographic comparison of native and exotic plants.

In natural ecosystems, plants face multiple herbivore attacks and are under selection to optimize their defenses to maximize their fitness (Lankau & Strauss, 2008; Wise & Rausher, 2013; Poelman & Kessler, 2016). We found that different latitudinal gradient of herbivory attributed to different herbivore feeding guilds (Figure 2a), which is consistent with the results of previous study (Zvereva & Kozlov, 2021). We also found that significant latitudinal gradient in herbivory driven by several major components of the herbivore feeding guild only occurs in native plants, but not in exotic plants (Figure 2b). Typically, to cope with a range of attackers that may all require different defensive traits, native plants have evolved mechanisms to recognize the specific guild of attackers by their feeding pattern, feeding position and induced defense phenotype against the specific attacking herbivore (Züst & Agrawal, 2016; Karban, 2019). However, exotic plants need to re-establish associations with multiple herbivore guilds after introduced into novel environments, a long-term co-evolutionary process (Joshi & Vrieling, 2005; Parker et al., 2006). Therefore, we can infer that the differences in the ability to respond to herbivore feeding guilds, result in different intensities of latitudinal gradients in herbivory between native and exotic plants. Our results provide a potential reason for the homogenized latitudinal gradient of herbivory observed in exotic plants.

Climatic factors have emerged as a pivotal selective pressure in the co-evolutionary processes driving the herbivory (Hamann et al., 2020). Moderately elevated temperatures can increase plant growth (Grace, 1987), productivity and secondary metabolite synthesis (Rustad et al., 2001; Zvereva & Kozlov, 2006; Pincebourde et al., 2017), and can also enhance the metabolism of heterothermic herbivores (Bale et al., 2002), thereby promoting increased consumption of plant tissues (Cornelissen, 2011; Jamieson et al., 2012). Reduced precipitation has been shown to increase plant chemical defenses and reduce herbivore performance due to limited access to nutrients (Gely et al., 2019). Our results showed that the positive significant correlations between climatic factors effect sizes (including BIO1, BIO12, BIO13 and BIO14) and herbivory effect sizes of native plants, while BIO4 effect sizes exhibited an opposite gradient (Figure 3). For exotic plants, we found no significant relationships in exotic plants with climatic factors (Figure 3). Thus, greater spatial heterogeneity in temperature and precipitation along the latitude are important predictors of the latitudinal pattern of herbivory only for native plants, but not for exotic plants. We also demonstrated that larger study spans are also an important predictor of the intensity of latitudinal gradients in herbivory (Figure S2). This may be due to the fact that studies spanning a larger area have greater climatic variability, which would result in a more significant latitudinal herbivory gradient. Therefore, expanding the study area in future studies may help researchers discover more significant biogeographic patterns. In short, these intricate relationships we found between climate and herbivory in native plants may have undergone long-term co-evolution, but this process is clearly lacking in exotic plants.

Our results showed that the latitudinal herbivory pattern of native plants observed in the field exhibited a significant latitudinal gradient in intensity, whereas that observed in common gardens did not (Figure S3). We did not observe significant latitudinal herbivory patterns of exotic plants in common gardens and field surveys. The comparison of differences in latitudinal herbivory gradients between the field and the common garden might indicate the response ability of herbivory-related traits in both plant identities to different environmental gradients (i.e., phenotypic plasticity; Guo et al. unpublished paper). Phenotypic plasticity is an inherent ability of plants to respond to highly heterogeneous environments by producing certain phenotypic changes in adaptive novel environments, as well as during evolution (Figure S3; West-Eberhard 1989). Long-term phenotypic plasticity results in plants that have the highest proportion of optimal phenotypes to adapt local herbivory and pass these phenotypes on to future generations through genetic differentiation during long-term coevolution (Rasmann et al., 2012). Our result implies the ‘adaptive plasticity’ in the latitudinal gradient pattern of herbivory in native plants, which is an active response of plants to environmental changes and is a heritable variation that has developed in plants over long-term evolutionary periods (Figure S4; Bhattarai et al., 2017). Conversely, for exotic plants, no significant latitudinal gradients of herbivory were observed in the field or in the common garden. This suggested that exotic plants tend to maintain homeostasis in highly heterogeneous environments, possibly driven by ‘generalist genotypes’ (Richards et al., 2006). However, it remains plausible that no any significant gradients in herbivory were observed for exotic plants, because they have not yet adapted to environment in new ranges.

We also described the intensity of latitudinal gradient in herbivory of exotic plants in relation to the time of introduction in our results, and found that exotic plants tended to show a significant increase in herbivory effect sizes with increasing time of introduction (Figure S3). This result suggests that longer time of introduction time may allow exotic plants to gradually adapt to multiple climatic factors and herbivore feeding guilds over a larger area of the introduction area, gradually producing different genotypes through physiological mechanisms such as phenotypic plasticity, which in turn promotes stronger patterns of latitudinal gradients (Molina-Montenegro & Naya, 2012). Accordingly, we can infer that the co-evolution of exotic plants with the novel environment of the introduced area over a long temporal scale contributes to generate the observed intensity of the latitudinal gradient.

As frequently encountered in ecological and evolutionary biology studies, the effect sizes exhibited heterogeneity across the dataset (Senior et al., 2016), which mainly between studies, may be due to differences in study design (e.g., differences in study scale, observation time and ecosystem) and differences between plant species. In our study, we controlled the latitude effect as the main factor, and limited other factors, so as to avoid the bias caused by different study designs as much as possible. Thus, we did not find the apparent bias of effect sizes across the dataset (Figure 4a). We also did not find that the journal impact factors and the publication time of studies had an effect on the effect sizes (Figure 4b, c). These results suggest that the quality of the journal and the publication time of the article did not bias the effect sizes in our dataset. In addition, our database focuses on studies of invasive and native plants, but there are currently a relatively few studies on the latitudinal gradient of herbivory of invasive plants, which may again introduce some bias.

Worldwide, more than 10,000 exotic plant species have now been introduced from their original range to colonies in new environments (van Kleunen et al., 2015). A number of these plants can gradually expand their distribution and adapt to the novel environment, thereby occupying the niches of local native plants and causing serious ecological damage to the introduced ecosystem (Christina et al., 2020). For example, *Spartina alterniflora*, *Ambrosia artemisiifolia*, and *Solidago canadensis*, these widely exotic plants have a high invasiveness and adaptability due to phenotypic plasticity and often evolve towards increased competitiveness in their introduced range (An et al., 2009; Sun & Roderick, 2019; Liu et al., 2020; Cheng et al., 2021). The spatial heterogeneity in latitudinal pattern of herbivory that we found, may reflect the unequal selection pressure between native and exotic plants (mainly invasive plants), which may directly or indirectly promote the rapid spread of invasive plants in their introduced area and facilitate the success of large-scale plant invasions (Huang et al., 2012; Schultheis & MacGuigan, 2018). Therefore, a better understanding of the biogeographic differences in herbivory between native and invasive plants, especially in the early stages of their introduction, may be essential (Harvey et al., 2010; Bhattarai et al., 2017). In addition, the relatively stable relationship between native plant herbivory and biotic and abiotic factors may be gradually disrupted as global climate change intensifies, while exotic plants may also experience climate change effects as they adapt to novel environments (Lu et al., 2015; Welshofer et al., 2018). These are likely to cause rapid fluctuations in the latitudinal patterns of plant herbivory. Therefore, future studies should include various factors such as global climate change, when examining the biogeographic pattern in herbivory.

## Supporting Information

**Table S1.** The restricted maximum likelihood (REML) models for full dataset and independent moderator variables. Moderator variables whose 95% confidence intervals (CI) do not cross zero are shown in bold; LCI 95% lower CI, UCI 95% CI; effect sizes (*z_r_*) are the response variable for this REML model.

| Variable | Estimate | SE | *z* | *p* | LCI | UCI |
| --- | --- | --- | --- | --- | --- | --- |
| *Full dataset* | **-0.127** | 0.054 | -2.366 | 0.018 | -0.231 | -0.022 |
| *Plant identities* |  |  |  |  |  |  |
| Native plants | **-0.130** | 0.054 | -2.417 | 0.016 | -0.235 | -0.025 |
| Exotic plants | -0.108 | 0.066 | -1.643 | 0.101 | -0.236 | -0.021 |
| *Observation types* |  |  |  |  |  |  |
| Field survey | **-0.130** | 0.053 | -2.442 | 0.015 | -0.235 | -0.026 |
| Common garden | -0.059 | 0.064 | -0.932 | 0.352 | -0.184 | 0.065 |
| *Herbivore feeding guilds* |  |  |  |  |  |  |
| All folivores | **-0.127** | 0.057 | 2.242 | -0.025 | -0.238 | -0.016 |
| Defoliators | **-0.130** | 0.057 | -2.295 | 0.022 | -0.241 | -0.019 |
| Gallers | -0.040 | 0.065 | -0.621 | 0.534 | -0.168 | 0.087 |
| Gazers | -0.376 | 0.500 | -0.752 | 0.452 | -1.355 | 0.604 |
| Miners | -0.056 | 0.065 | -0.863 | 0.388 | -0.183 | 0.071 |
| Sap feeders | **-0.171** | 0.066 | -2.569 | 0.010 | -0.301 | -0.040 |
| Seed feeders | **-0.215** | 0.092 | -2.329 | 0.020 | -0.396 | -0.034 |
| Stem feeders | **-0.349** | 0.085 | -4.107 | <.000 | -0.515 | -0.182 |

**Table S2.** The restricted maximum likelihood (REML) model result for the interaction between plant identities (native/exotic) and herbivore feeding guilds. Moderator variables whose 95% confidence intervals (CI) do not cross zero are shown in bold; LCI 95% lower CI, UCI 95% CI; Effect sizes (*z_r_*) is the response variable for this REML model.

| Variable | Estimate | SE | *z* | *p* | LCI | UCI |
| --- | --- | --- | --- | --- | --- | --- |
| Native × all folivores | **-0.138** | 0.058 | -2.396 | 0.017 | -0.250 | -0.025 |
| Native × defoliators | **-0.148** | 0.058 | -2.565 | 0.010 | -0.261 | -0.035 |
| Native × gallers | -0.059 | 0.066 | -0.897 | 0.370 | -0.188 | 0.070 |
| Native × gazer | -0.376 | 0.499 | -0.754 | 0.451 | -1.353 | 0.601 |
| Native × miners | -0.087 | 0.067 | -1.299 | 0.194 | -0.217 | 0.044 |
| Native × sap-feeders | -0.105 | 0.070 | -1.515 | 0.130 | -0.242 | 0.031 |
| Native × seed-feeders | -0.095 | 0.102 | -0.933 | 0.351 | -0.295 | 0.105 |
| Native × stem-feeders | **-0.427** | 0.089 | -4.796 | <.0001 | -0.602 | -0.253 |
| Exotic × all folivores | -0.079 | 0.074 | -1.066 | 0.286 | -0.224 | 0.066 |
| Exotic × defoliators | -0.061 | 0.083 | -0.736 | 0.462 | -0.223 | 0.101 |
| Exotic × gallers | 0.1594 | 0.173 | 0.921 | 0.357 | -0.180 | 0.499 |
| Exotic × miners | 0.121 | 0.137 | 0.882 | 0.378 | -0.148 | 0.390 |
| Exotic × sap-feeders | **-0.279** | 0.089 | -3.137 | 0.002 | -0.454 | -0.105 |
| Exotic × seed-feeders | **-0.512** | 0.176 | -2.915 | 0.004 | -0.857 | -0.168 |
| Exotic × stem-feeders | 0.092 | 0.177 | 0.518 | 0.604 | -0.255 | 0.438 |

**Table S3.** The restricted maximum likelihood (REML) model result for the interaction between plant identities (native/exotic) and observation types (field survey/common garden). Moderator variables whose 95% confidence intervals (CI) do not cross zero are shown in bold; LCI 95% lower CI, UCI 95% CI; Effect sizes (*z_r_*) is the response variable for this REML model.

| Variable | Estimate | SE | *z* | *p* | LCI | UCI |
| --- | --- | --- | --- | --- | --- | --- |
| Native × field | -0.119 | 0.081 | -1.471 | 0.141 | -0.278 | 0.040 |
| Native × garden | -0.072 | 0.068 | -1.051 | 0.293 | -0.203 | 0.062 |
| Exotic × field | -0.033 | 0.065 | -0.502 | 0.616 | -0.160 | 0.095 |
| Exotic × garden | -0.141 | 0.053 | -2.656 | 0.008 | -0.246 | -0.037 |

**Figure S1.**
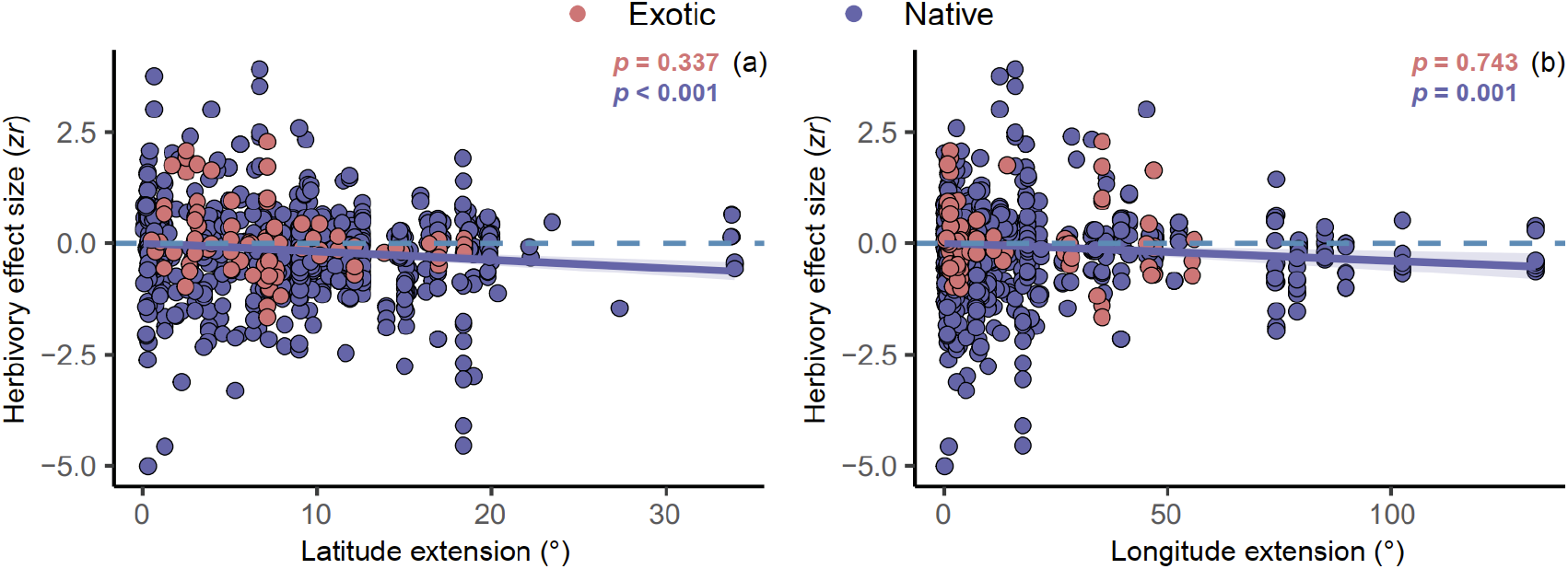
The relationships between effect sizes (*z*_r_) and latitudinal extension (a), and longitudinal extension (b). For each panel, *p* < 0.05 indicates a significant relationship. The envelopes represent the 95% CI of the linear regression.

**Figure S2.**
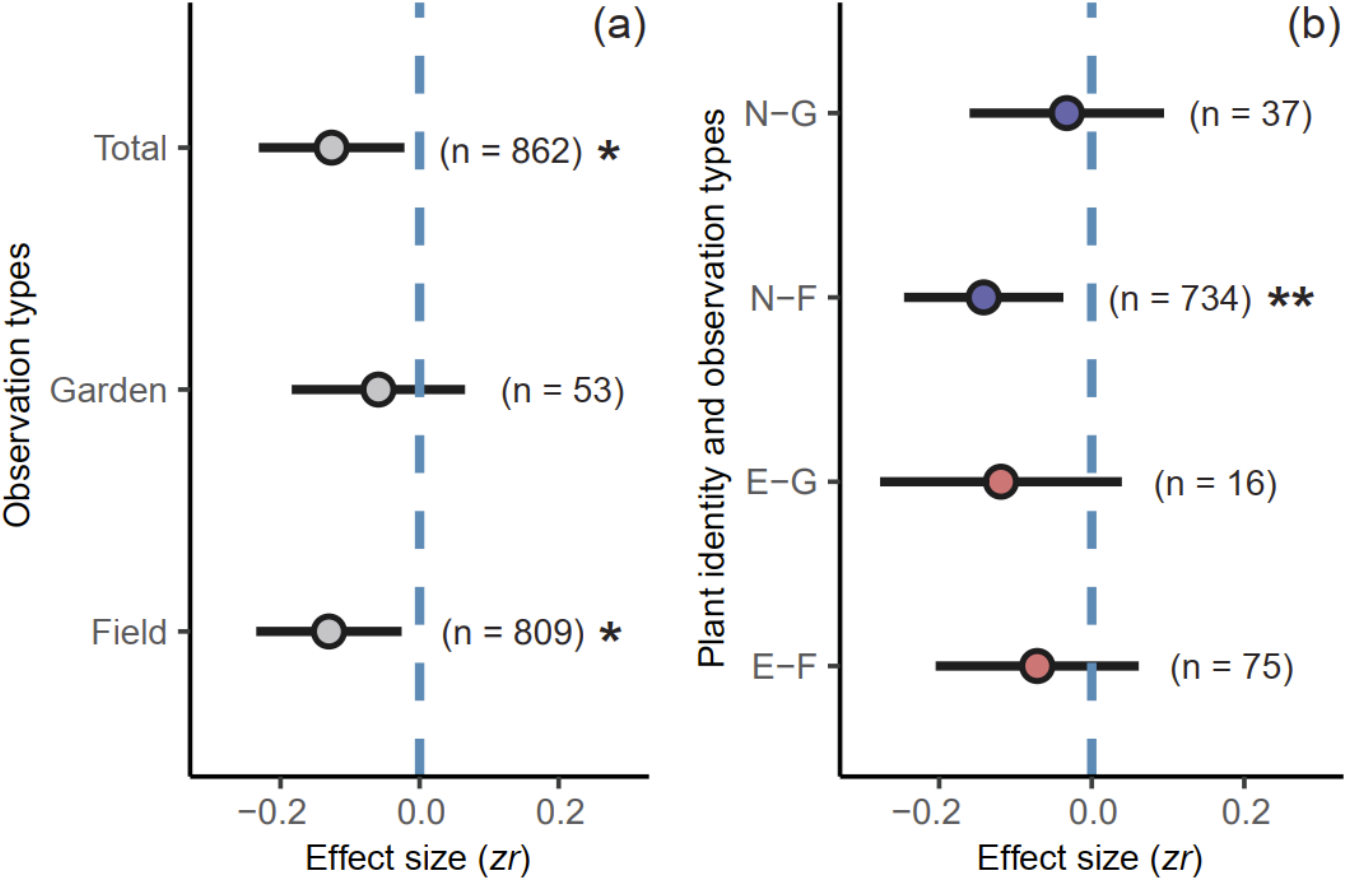
The role of plant identities and observation types on effect sizes. On the panel (a), Garden and Field mean the effect sizes were observed from common garden experiments and field surveys, respectively. On the panel (b), N and E indicate the abbreviation of two level in plant identity (i.e., native and exotic). G and F indicate the abbreviation of two level in observation types (i.e., common garden and field survey). On the graph, the negative effect sizes indicate a decreased interaction intensity with increased latitude. The estimates and 95% confidence intervals (i.e., horizontal lines) presented here are from restricted maximum likelihood (REML) models. Numbers of effect sizes are shown in parentheses. Statistical significance is indicated as: *, *p* < 0.05; **, *p* < 0.01; ***, *p* < 0.001.

**Figure S3.**
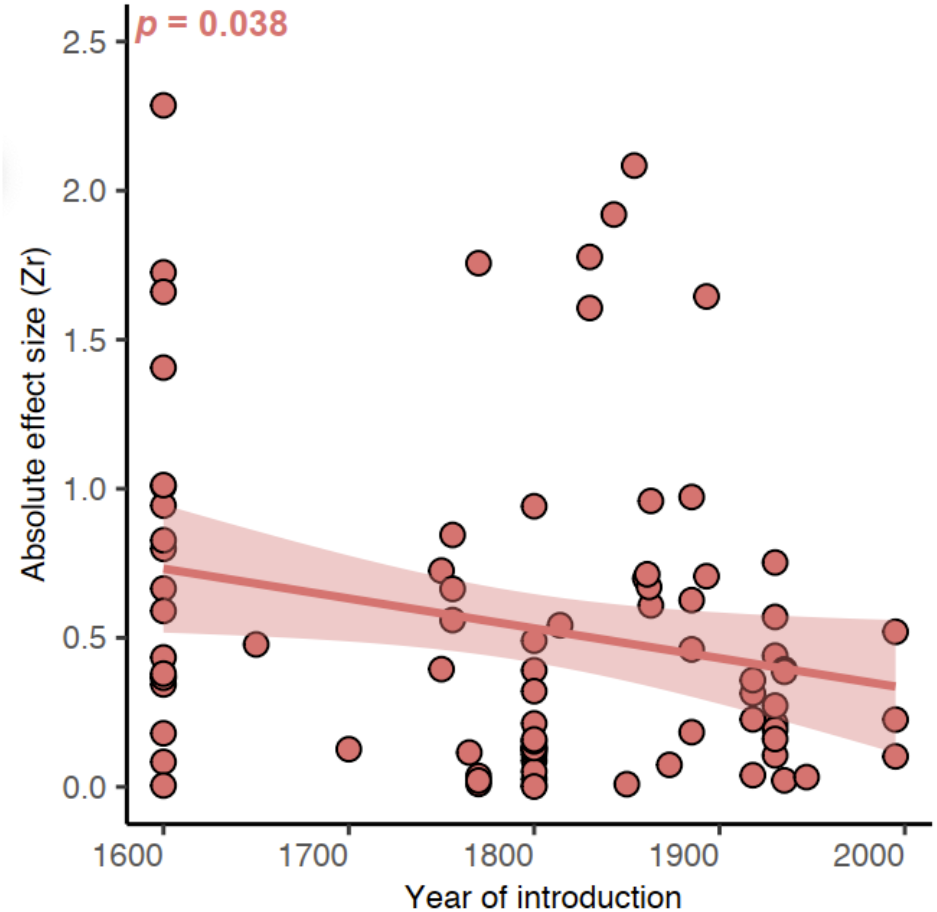
The relationship between of year of introduction and absolute effect sizes (*z*_r_). On each panel, *p* < 0.05 indicates a significant relationship. The envelopes represent the 95% CI of the linear regression.

**Figure S4.**
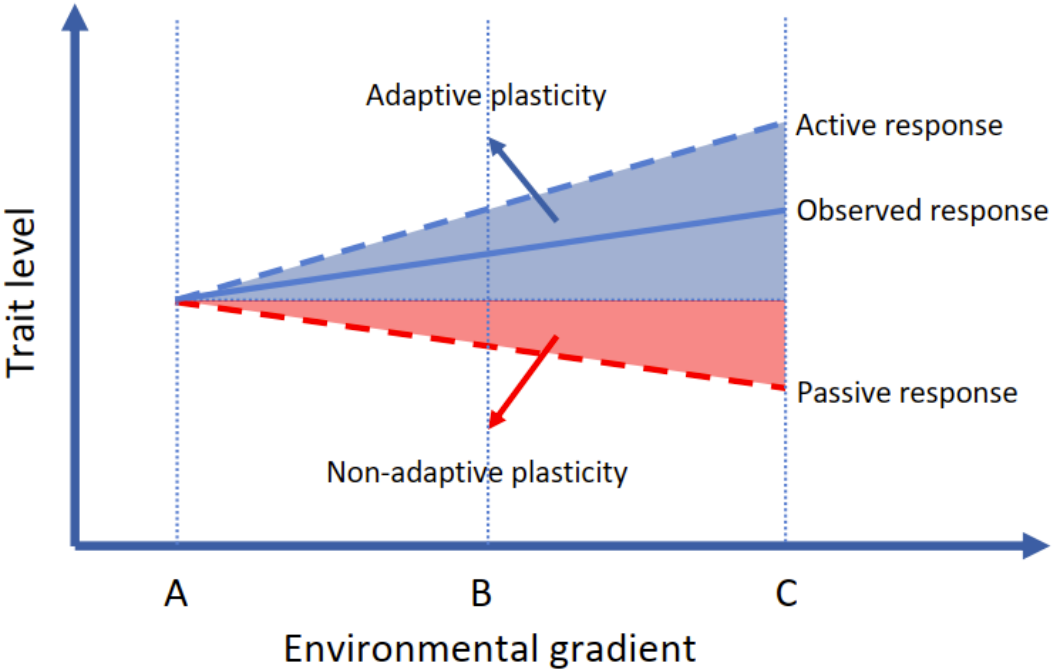
Conceptual diagram of the relationship between environmental gradient and traits level. (refers to van Kleunen & Fischer, 2005)

